# Ketamine strengthens synaptic inputs to the dorsal raphé to boost serotonergic activity: pivotal for rapid antidepressant-like effects

**DOI:** 10.1101/2025.10.24.684353

**Authors:** Soukaina Es-Safi, Céline Bourcier-Lucas, Erika Abrial, Adeline Cathala, Thierry Lesté-Lasserre, Philippe De Deurwaerdère, Jean-Michel Revest, Jean Mazella, Nasser Haddjeri, Guillaume Lucas

## Abstract

Ketamine at subanaesthetic dose is known as a fast-acting antidepressant (AD), able to facilitate synaptic plasticity in the medial prefrontal cortex (mPFC) or the hippocampus. However, its influence on the serotonergic (5-HT) system is more confusing as it loses its behavioral effects in 5-HT-depleted animals, but does not modify 5-HT neuron function. We hypothesized that this discrepancy was due to the different temporal scales chosen in the related studies. We performed electrophysiological recordings of 5-HT neurons in the dorsal raphé nucleus (DRN) and microdialysis measurements of 5-HT release in the ventral hippocampus of male Sprague-Dawley rats. Experiments were designed to collect the results over a long duration, i.e. 4-5 hours after the injection. Levels of the pro-neuroplastic factors PSD-95 and Synapsin-1 in the DRN were also assessed, as was cell proliferation in the dentate gyrus (DG). 5-HT neuron mean firing rate was unmodified within the 2 h that followed ketamine (10 mg/kg, i.p.), but strongly (90%) increased in the 2-5 h time bin, an effect abolished by lesioning the mPFC or administering the mTOR inhibitor Torin-2. A similar kinetics was found for hippocampal 5-HT release. Synapsin-1 and PSD-95 mRNA transcriptions were enhanced at 2 h, and PSD-95 protein levels appeared to peak at 24 h. Finally, DG mitogenesis was enhanced 48 h post-injection, an augmentation suppressed in 5-HT-depleted animals. Low-dose ketamine provokes a “neuroplastic wave” originating in the mPFC and emerging in the hippocampus, a transfer in which the 5-HT system appears to act as an integrative hub of plasticity.

**SIGNIFICANCE STATEMENT:** This study helps reconcile two theories, often opposed to explain antidepressant (AD) action: the “serotonergic hypothesis” proposing that AD efficacy primarily results from an increase of serotonin, and the “neuroplastic theory” whereby only significant changes in brain connectivity can explain mood improvement. Here we show that ketamine, a drug exerting AD effects much faster than classical molecules do, augments synaptic strength onto serotonergic neurons, which subsequently triggers an elevation of serotonin. In turn, this enhanced serotonergic neurotransmission leads to the production of new neurons in the hippocampus, therefore reshaping the circuitry of this brain area. The serotonergic system appears to transfer the neuroplastic changes induced by ketamine across the brain, thus behaving like an integrative hub for its AD action.

## INTRODUCTION

For almost two decades now, the use of the non-selective N-methyl-D-aspartic acid (NMDA) receptor antagonist ketamine at subanaesthetic doses has proved effective as an antidepressant (AD) treatment in clinical use (1–4). More importantly, it has been highlighted as a fast-acting AD (1,3,5), a crucial feature in regard to the major challenge that constitutes the delayed efficacy of canonical molecules (6–7). This qualification of “fast-acting” however, needs to be further precised, as it actually covers different temporal realities. Indeed, some AD effects of ketamine have been described within timeframes that can be qualified as “immediate”, i.e. a few tens of minutes only (< 2 hours) after its administration (1). Other reports have stressed its ability to display AD efficacy after one or two days, corresponding to what is generally expected as a “rapid” onset of AD action (2,5,8–9). Similarly, in rodent models specifically designed to take into account the kinetics of AD treatments such as the chronic mild stress or the learned helplessness, a low dose (10 mg/kg) of ketamine is able to rapidly (24 h) reverse the depressed-like phenotype (10–12). Regarding the mechanisms involved, several studies have identified the medial prefrontal cortex (mPFC) as a key brain area, for both the immediate, and rapid time scales. Thus, within minutes after its systemic administration, ketamine induces an increase of glutamate release and metabolic cycling in the mPFC, effects that are accompanied by an activation of synaptic signaling protein pathways in the same structure (13–15). Also, a single administration of ketamine at a subanaesthetic dose is sufficient to produce synaptogenesis within 1-2 days in the mPFC of rodents, either in resting conditions or in a model of depression (11,13,16). Together, these data further strengthen the idea that AD efficacy results from a restored and/or improved connectivity between brain neuronal networks, a concept that has emerged about twenty years ago as the “plastic theory” of depression (17–19).

The fact remains though, that numerous evidences have also pointed out the central role played by monoamines, and more specifically the serotonergic (5-HT) system of neurotransmission, in the AD mechanisms of action. The “5-HT hypothesis” of depression was one of the firsts proposed to address the question of AD effectiveness (for review, 6), and is still put forward nowadays, including in the search for a rapid onset of action (20–21). And indeed, several behavioral studies have shown that the AD-like effects of ketamine are suppressed after a brain depletion of 5-HT in both rats and mice, implying that this neurotransmitter is essential for their implementation (22–26). However, researches aimed at addressing the influence of ketamine on the 5-HT system at the functional level led to conflicting results. Thus, the activity of 5-HT neurons within the dorsal raphe nucleus (DRN), the area which harbors the vast majority of brain 5-HT cell bodies (27), was found to be either unaffected in rats (20,28), or even decreased in mice (26) after the administration of low doses of ketamine. One possibility to account for these discrepancies may precisely be related to the different time scales chosen to perform those studies. In rats, all behavioral data obtained within a rapid timeframe (i.e. 24-48 h after the injection) show a 5-HT dependency for the AD-like effects of ketamine (22,24–25), a property which is not found in the immediate (60 min) scale of time (22). Now, most of the functional (electrophysiological and neurochemical) 5-HT neuron recordings currently available have been specifically designed to fit within a period of less than 2 hours (20, 28–29). The question that can be raised is therefore the following: what if the involvement of central 5-HT neurons in the antidepressant-like properties of ketamine would itself result from processes of increased neuroplasticity? Indeed, the stimulation of neurons projecting from the mPFC to the DRN is able to trigger AD-like effects through the mobilization of endogenous 5-HT (30–32). Considering that the activation of the mPFC by ketamine appears to be already dependent on synaptogenesis (11,13), it cannot be ruled out that similarly, the ensuing facilitation of DRN 5-HT activity also requires synaptic growth to manifest. This would account for the apparent “delayed” intervention of 5-HT in the results mentioned above.

The present study was aimed at addressing this hypothesis. To this purpose, electrophysiological and neurochemical experiments were designed in rats, to follow the activity of 5-HT neurons several hours after the administration of a subanaesthetic dose of ketamine. In parallel, the levels of the molecular markers of synaptic plasticity PSD-95 and synapsin I were measured in the DRN 2, 6 and 24 hours after ketamine. Finally, we also assessed the effect of ketamine on cell proliferation in the dentate gyrus (DG) of the hippocampus, considered a hallmark of AD efficacy (33), in the presence or absence of endogenous 5-HT 48 hours after the injection.

## RESULTS

### Effect of ketamine on the firing activity of DRN 5-HT neurons

In agreement with our earlier results (34), the mean firing rate of DRN 5-HT neurons in vehicle-treated rats was found to be around 1.3 Hz (**Fig. 1A**). Interestingly, this value did not appear to be affected in the particular conditions of long duration chloral hydrate anaesthesia we used; as shown in **Fig. 1B**, even after more than 4 hours, the neurons encountered along recording tracks still discharged within the 1-1.5 Hz range. By contrast, in the same conditions 5-HT neuronal activity reached 2.45 Hz on average, when recorded within a 2-5 hours timeframe after the administration of ketamine at the dose of 10 mg/kg, i.p. This corresponds to an 90% increase, a statistically significant effect (Student’s t-test, t_9_ = −4.32, p < 0.01, **Fig. 1A**). As shown in **Fig. 1B**, it was not uncommon to encounter neurons discharging at 4, or even close to 5 Hz, in ketamine-injected animals.

**Figure 1.**
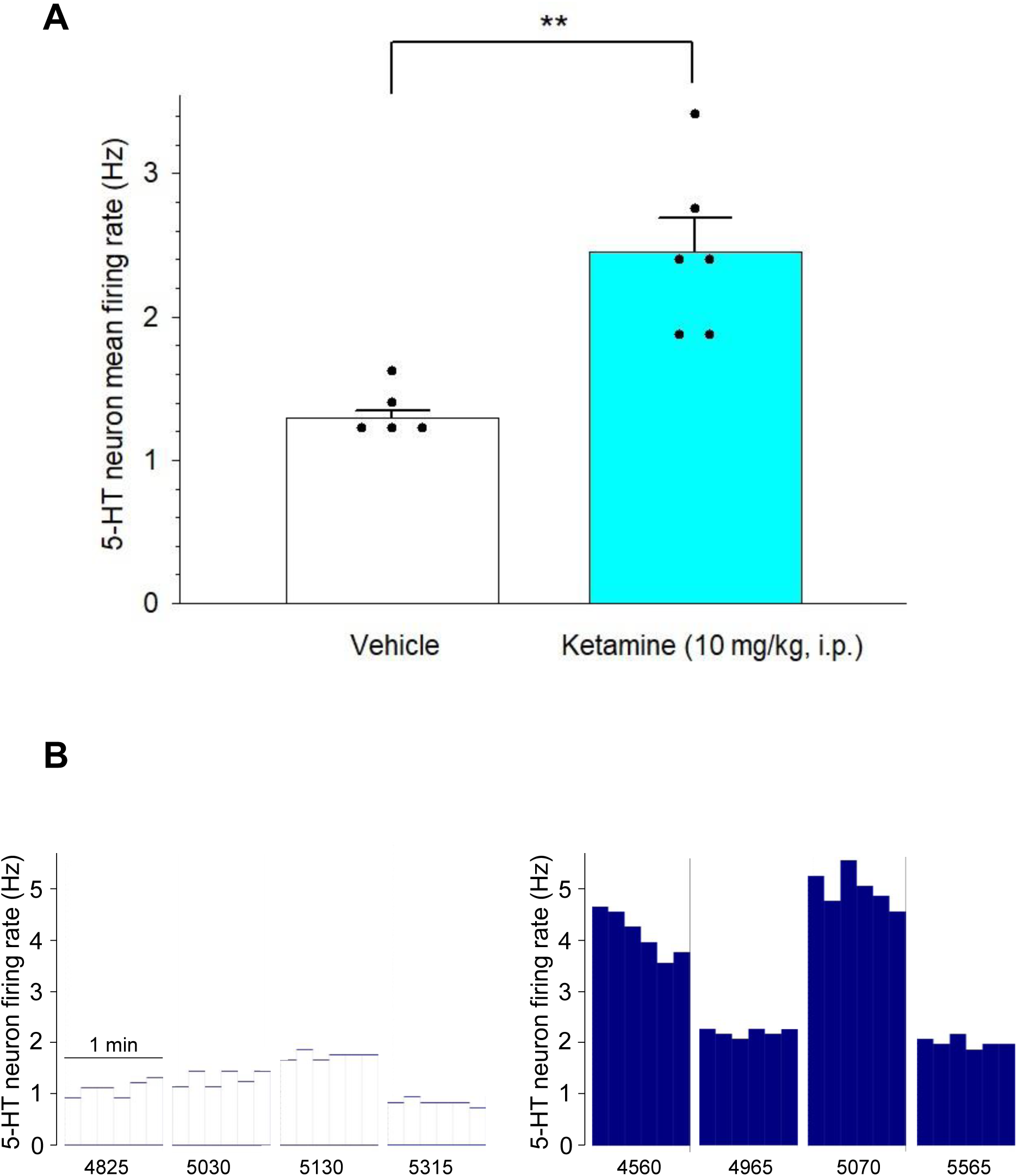
Effect of ketamine on the firing activity of DRN 5-HT neurons. Ketamine (10 mg/kg) or its vehicle was administered intraperitoneally. Recordings started at least 120 min after the injection, and were performed for a maximal duration of 300 min thereafter. **A**: Each dot represents the average 5-HT firing activity of one single rat, calculated on the basis of 8-12 neurons recorded along successive tracks performed within the DRN. Bar histograms summarize the mean (± S.E.M.) of these individual data. ** p < 0.01 vs Vehicle, Student’s *t*-test. Total number of neurons recorded: n = 44 (Vehicle) and 62 (Ketamine). **B**: Integrated firing rate histograms showing representative samples of 5-HT neurons recorded from descents performed along the DRN in vehicle-(left panel) and ketamine (right panel)-treated rats, respectively. In both cases, the descent had been performed around 210-240 min after the injection. Each cluster of bars summarizes the electrical activity of one neuron, one bar representing the average number of recorded action potentials per 10 s. The numbers below the abscissa axis represent the height (in mm, and related to the cortical surface) at which each neuron was encountered, according to the micromanipulator system display.

### Effect of ketamine on 5-HT neurons in the presence of a lesion of the mPFC

Ketamine at subanaesthetic doses has been shown to enhance the activity of mPFC pyramidal neurons within minutes after its administration, a property which appears to be pivotal in its AD-like effects (13,15). Further, it is known that a stimulation of the mPFC is able to trigger the firing of DRN 5-HT neurons (30,32). Thus, we next tested the influence of ketamine, in the same conditions of recording and drug administration, but after an electrolytic lesion of the mPFC (36). As shown in **Fig. 2A**, and in agreement with previous results (34), the lesion had no effect in vehicle-treated rats, which displayed an average firing rate of 1.27 Hz. However, ketamine induced a slight, non-significant decrease of 5-HT activity in lesioned animals (Student’s *t*-test, t_7_ = 1.46, n.s., **Fig. 2A**), indicating that an electrolytic lesion of the mPFC actually abolishes the facilitatory control evidenced in **Fig. 1**.

**Figure 2.**
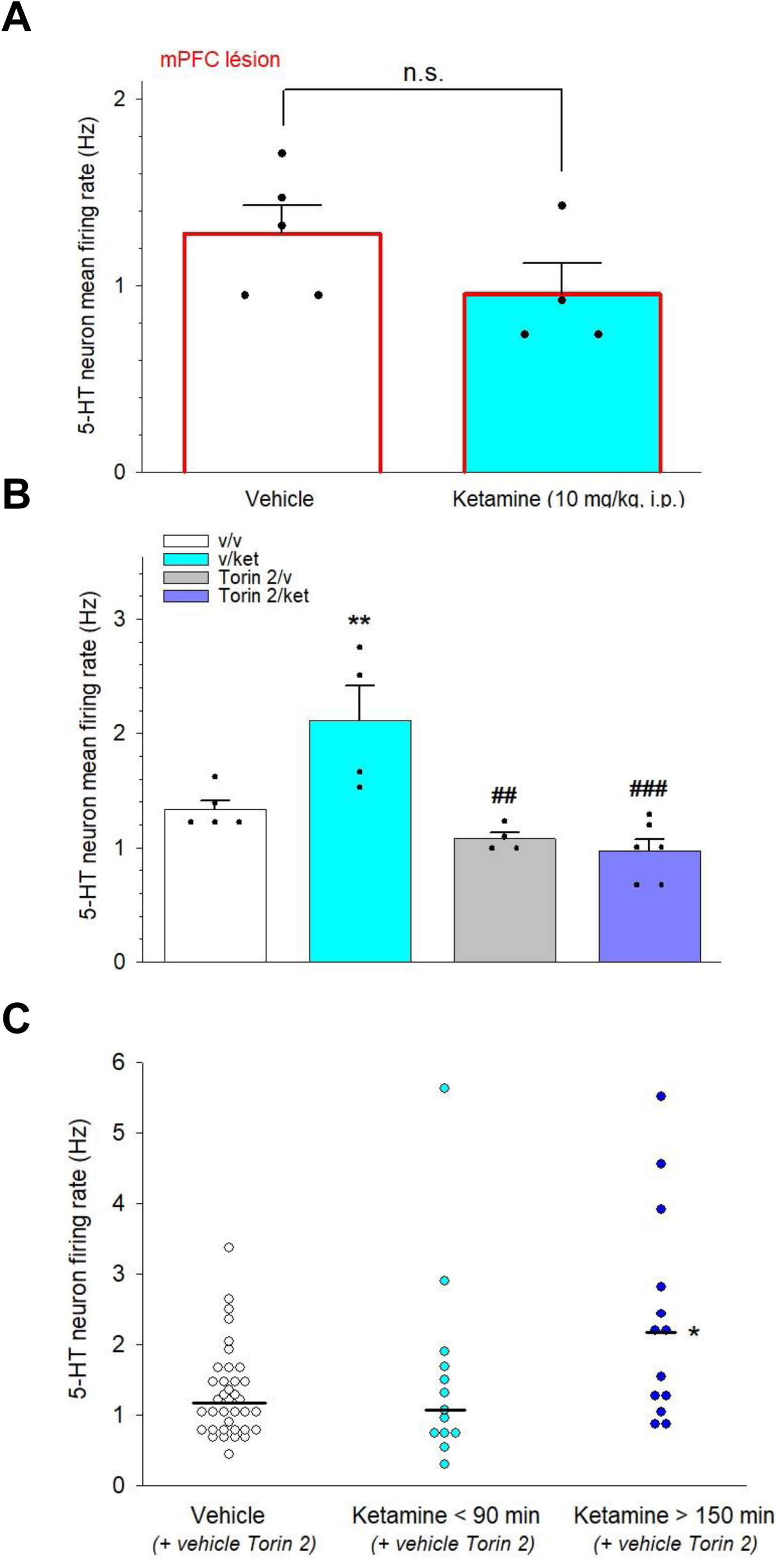
Involvement of the mPFC and of the mTOR signaling pathway in the effect of ketamine on 5-HT neuron activity. **A**: Comparison of the effects of ketamine (10 mg/kg, i.p.) and its vehicle in rats bearing an electrolytic lesion of the mPFC. The lesion (500 µA direct current delivered in the mPFC during 10 s) was made just before the surgery for electrophysiological recordings. Ketamine (10 mg/kg) or its vehicle was administered intraperitoneally, recordings started at least 120 min after the injection and were performed for a maximal duration of 300 min thereafter. Each dot represents the average 5-HT firing activity of one single rat, calculated on the basis of 8-12 neurons recorded along successive tracks performed within the DRN. Bar histograms summarize the mean (± S.E.M.) of these individual data. Total number of neurons recorded: n = 47 (Vehicle) and 38 (Ketamine). **B**: Effect of Ketamine (ket; 10 mg/kg, i.p.) and of the mTOR pathway inhibitor Torin 2 (1 mg/kg, i.p.) or their vehicles (v) on 5-HT neuron mean firing rate within the DRN. Recordings started 30 min after the injection of ketamine and lasted up to 240 min thereafter. Torin 2 or its vehicle was given 15 minutes before ketamine. Each dot represents the average 5-HT firing activity of one single rat, calculated on the basis of 7-10 neurons recorded along successive tracks performed within the DRN. Bar histograms summarize the mean (± S.E.M.) of these individual data. ** p < 0.01 vs v/v; ## p < 0.01, ### p < 0.001 vs v/ket, Newman-Keuls test after significant one-way ANOVA. Total number of neurons recorded: n = 37 (v/v), 40 (v/ket), 38 (Torin 2/v) and 52 (Torin 2/ket). **C**: Distribution of individual 5-HT neuron firing rates from the v/v and v/ket groups of panel **B**, in function of the post-injection time at which they have been recorded. In the v/ket group, the cells recorded between 90 and 120 min after the administration of ketamine were discarded, so that only those taken before 90 min or after 120 min have been plotted. Since no differences were observed between < 90 min, 90-120 and > 150 min values in vehicle-injected rats, they have been pooled together in the “Vehicle” column. Horizontal bars represent the median of each group. * p < 0.05 vs Vehicle, Dunn’s test after significant one-way ANOVA on ranks.

### Influence of Torin 2 on the facilitatory effect of ketamine

Several studies have reported that the ability of ketamine to mobilize mPFC pyramidal neurons involves an activation of the mammalian Target of Rapamycin (mTOR) signaling pathway (11–13,35). To assess whether this mechanism is also responsible for the observed facilitation of 5-HT activity, the effect of ketamine on DRN 5-HT neuron firing rate was tested in the presence of the highly selective mTOR inhibitor Torin 2 (36), at a dose (1 mg/kg i.p.) that has been reported to stand within the effective range *in vivo* (36–37). In these experiments however, we could not wait until 120 min after the administration of ketamine, as we did in the previous ones. Indeed, Torin 2 had to be administered beforehand, to make sure that the mTOR pathway would be already inhibited, at least partially, at the time ketamine would reach the brain. Because the half-life of Torin 2 is around 45 min in rodents (36), it was decided to give it 15 minutes before the subanaesthetic dose of ketamine, and to start the recordings 45 min thereafter (thus 30 min after ketamine). Still, recordings were continued until up to 4 hours post-injection of ketamine, to stay in the same long-term paradigm than in the above experiments. A classical four-group experimental design has been used (**Fig. 2B**). As revealed by a one-way ANOVA, a highly significant group effect was observed: F(3, 15) = 10.7, p < 0.001. More specifically, in the vehicle/ketamine group the values reached an average of 2.1 Hz, statistically different from those (1.33 Hz) found in the vehicle/vehicle one (Newman-Keuls *post-hoc* test, p < 0.01, **Fig. 2B**). It appears therefore that, in these experimental conditions, ketamine is still able to induce an increase of DRN 5-HT neuron mean firing rate. The Torin 2/vehicle and Torin 2/ketamine groups were significantly lower than vehicle/ketamine (Newman-Keuls test, p < 0.01 and p < 0.001, respectively), but more importantly, they were not different one from each other (**Fig. 2B**), showing that the facilitatory effect of ketamine is suppressed in the presence of Torin 2.

Interestingly, the increase observed after ketamine alone (2.1 compared to 1.33, thus +57%) was less prominent than in the first series of experiments. We hypothesized that this discrepancy was due to the different timings of drug administration and recordings. If, as proposed in the Introduction, a delay of ≈ 2 h is required before the activation of 5-HT neurons becomes observable, then the apparent dampening of ketamine effect would be due to the weight of the early-recorded neurons in the calculation of the mean. To test for this possibility, a statistical segregation was made: 5-HT neurons from the vehicle/ketamine group and recorded within 30-90 min after ketamine administration were pooled together, as well as those recorded more than 150 min thereafter; the “grey zone” corresponding to the 90-120 min timeframe was excluded. All the neurons from the vehicle/vehicle group have been kept together, as no temporal difference were expected, nor have been observed. Results are illustrated in **Fig. 2C**: because the distribution of individual firing rates did not follow a normal distribution, statistical comparisons were made by using non-parametric testing. A group effect was revealed when performing a one-way ANOVA on ranks (H_2_ = 6.7, p< 0.05), and the *post-hoc* Dunn’s test evidenced a significant difference between the “ketamine > 150 min” and vehicle (p < 0.05) medians (**Fig. 2C**). It is particularly striking that the value of the former (2.18 Hz) represented an increase of 86% when compared to the latter (1.17 Hz), a result very similar to the 90% augmentation reported in **Fig. 1A**. By contrast, the median of the “ketamine < 90 min” group (1.07 Hz) did not differ from that of the vehicle (Dunn’s test, n.s.). Altogether, these results confirm that a delay is necessary before an influence of ketamine can be observed on DRN 5-HT activity. Also, they may explain why no apparent effects were detected in previous studies, which recordings were done within the two hours that followed ketamine administration (28–29).

### Effect of ketamine on hippocampal 5-HT release

The hippocampus, and most notably its ventral part, appears to play a key role for the AD-like efficiency of ketamine (38). The next step of the study was therefore aimed at determining whether the activation of DRN 5-HT neuronal firing induced by ketamine translates into a mobilization of endogenous 5-HT in this particular area of interest. To this purpose, the technique of intracerebral microdialysis was used, in a long-lasting protocol as for electrophysiological experiments. In these conditions 5-HT extracellular levels slightly and progressively decreased over time in vehicle-treated rats, to reach around 70% of basal values at the end of the paradigm (**Fig. 3A**). It is likely that this effect was due to the anaesthetic chosen to perform microdialysis: indeed, isoflurane has been reported to exert an inhibitory influence on the impulse flow of some 5-HT neurons (39), and reciprocally activating 5-HT cell bodies promotes arousal from isoflurane anaesthesia (40). In our protocol, the stabilization period lasted 2 hours, and the microdialysis study in itself for an additional 4 hours, thus totaling 6 hours of exposure to the anaesthetic. It cannot be excluded that the accumulation of isoflurane during such a long period elicited changes in the homeostasis of the 5-HT system, leading to a progressive decline of its release. At any rate, ketamine at the dose of 10 mg/kg, i.p. was effective to modify this picture. Thus, while 5-HT extracellular levels remained unchanged during the first 100 min after the injection, they were statistically increased at t = 120 and t = 140 min (Student’s *t*-test, t_8_ = −3.72 and −2.34, p < 0.01 and p < 0.05 vs the same timepoint in the vehicle group, respectively; **Fig. 3A**). Although the Student’s *t*-tests were not significant afterwards, the kinetics clearly shows that in the ketamine group hippocampal 5-HT release stabilized around 110% of baseline until the end of the experiment (**Fig. 3A**). Averaging values over the entire post-injection period revealed a significant difference between the two groups (100% vs 77%, Student’s *t*-test t_9_ = −2.7, p < 0.05; **Fig. 3B**). Here again, the difference was due to the second half (120-240 min) of the period (108% vs 70%, Student’s *t*-test t_8_ = −2.4, p < 0.05), the means being almost identical during the first one (91% vs 86%, Student’s *t*-test t_9_ = −0.7, n.s.) (**Fig. 3B**). Thus, in striking similarity with electrophysiological experiments, our results show that ketamine is actually able to facilitate 5-HT outflow in the ventral hippocampus, but requires a delay of ≈ 2 h to do so.

**Figure 3.**
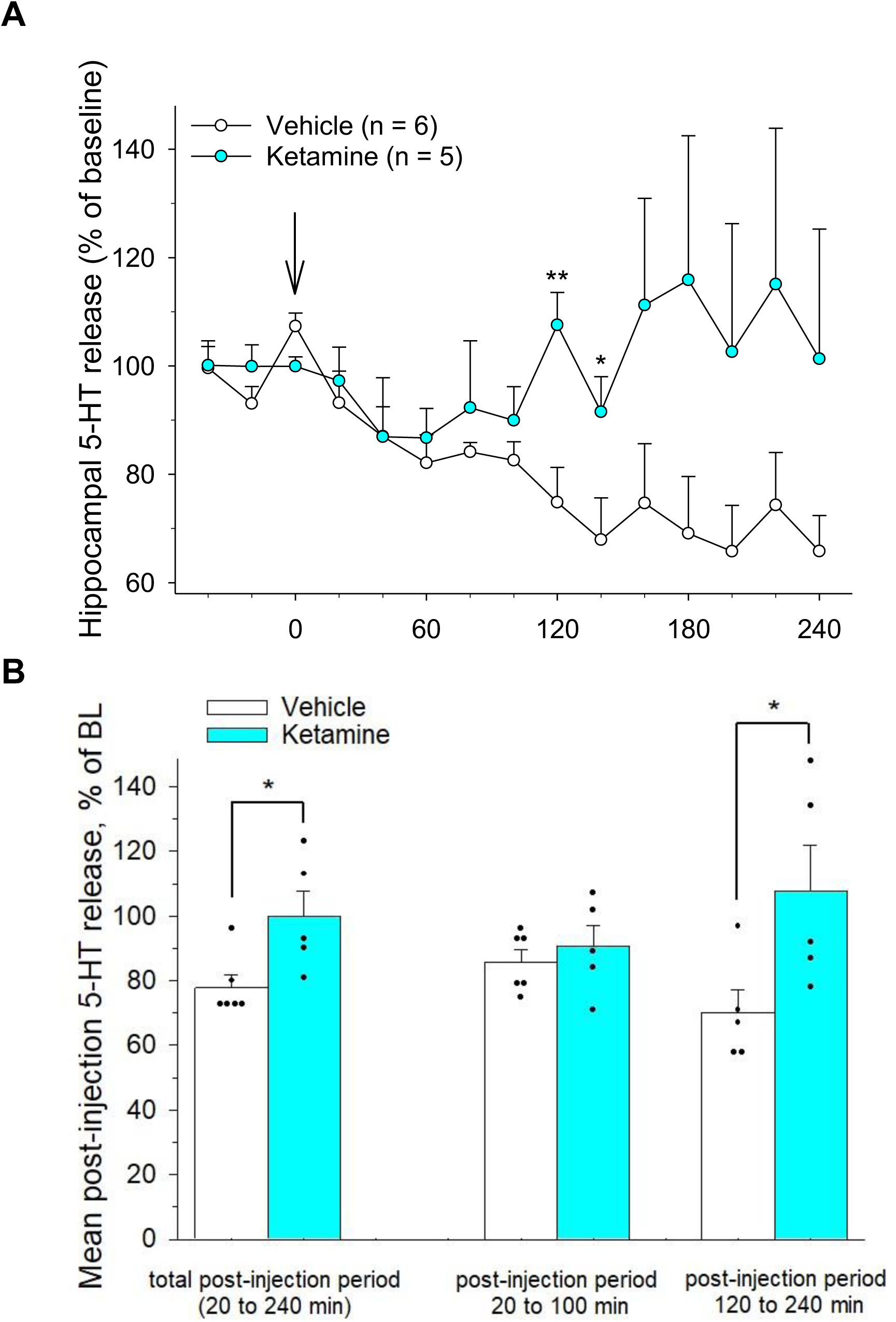
Effect of ketamine (10 mg/kg, i.p.) on the release of 5-HT within the ventral hippocampus, as measured by using *in vivo* intracerebral microdialysis. **A**: Time-course of 5-HT extracellular levels in vehicle-and ketamine-treated animals. Dialysate samples were collected every 20 minutes, and results are expressed as mean (± S.E.M.) percentage effect with respect to the baseline, calculated from the three timepoints preceding drug administration. As indicated by the *arrow*, ketamine was injected at the end of the third sample, chosen as time zero of the kinetics. * p < 0.05, ** p < 0.01 vs the same timepoint in the Vehicle group, Student’s *t*-test. **B**: Comparison of the average post-injection 5-HT release values in the two groups of panel **A**. Each dot represents 5-HT release, taken as percentage of baseline, averaged in one single rat over either the total post-injection period (timepoints 20 to 240 min, left), or in the early (timepoints 20-100 min, center) and late (timepoints 120-140 min, right) post-injection phases. In the vehicle group, one rat was lost after 100 min due to the detachment of one tubing from the canula. Bar histograms summarize the mean (± S.E.M.) of these individual data. * p < 0.05 vs Vehicle, Student’s *t*-test.

### Effect of ketamine on molecular markers of synaptic plasticity within the DRN

Ketamine is able to induce various changes in the brain within minutes after its administration (13–15). It appears therefore very unlikely that the delayed effects evidenced above would be due to pharmacokinetic factors. Rather, we hypothesized that the administration of a low dose of ketamine provokes a progressive synaptic strengthening in the DRN, as it has been shown to do in the mPFC (13,15). The levels of Synapsin I and PSD-95, respectively pre- and post-synaptic markers of increased plasticity (41–42) were thus measured in the DRN at different times after a single injection of ketamine (10 mg/kg, i.p.). Using the technique of RT-PCR, we found that the mRNA expression of both PSD-95 and Synapsin I was augmented by ≈ two-fold after 2 hours, a transitory effect no more observed at 6 and 24 hours post-administration (**Figs. 4A** and **4B**). In each case, a group effect was revealed by the one-way ANOVA [F(3, 17) = 4.83, p < 0.05 and F(3, 17) = 10.78, p < 0.001, respectively], with the “ketamine 2h” group being statistically different from the control (Dunnett’s test, p < 0.05 and p < 0.01, respectively; **Figs. 4A** and **4 B**). We next focused more specifically on the levels of the PSD-95 protein, which play a fundamental role in scaffolding and stabilizing postsynaptic densities, as well as the cohesion of NMDA and AMPA glutamatergic receptors during the formation of excitatory synapses (42). To this purpose, Western blots were performed in the same conditions of drug administration and brain extraction. The obtained results indicate that the protein expression increased over time, attaining its maximal value 24 h after the injection (**Figs. 4C** and **4D**). In these experiments, the one-way ANOVA failed to reach statistical significance [F(3, 20) = 3, p = 0.055], maybe because the gradual nature of the increase did not permit to make a group stand out above the others in this paradigm. However, by using paired comparisons, it was clear that protein levels at t = 24 h were significantly higher than in control animals (Student’s *t*-test, t_10_ = −3, p < 0.05). It appears therefore that ketamine at the dose of 10 mg/kg, i.p. is actually able to mobilize some key players of the synaptogenesis process within the DRN, within a timeframe compatible with the kinetics of its influence on 5-HT activity.

**Figure 4.**
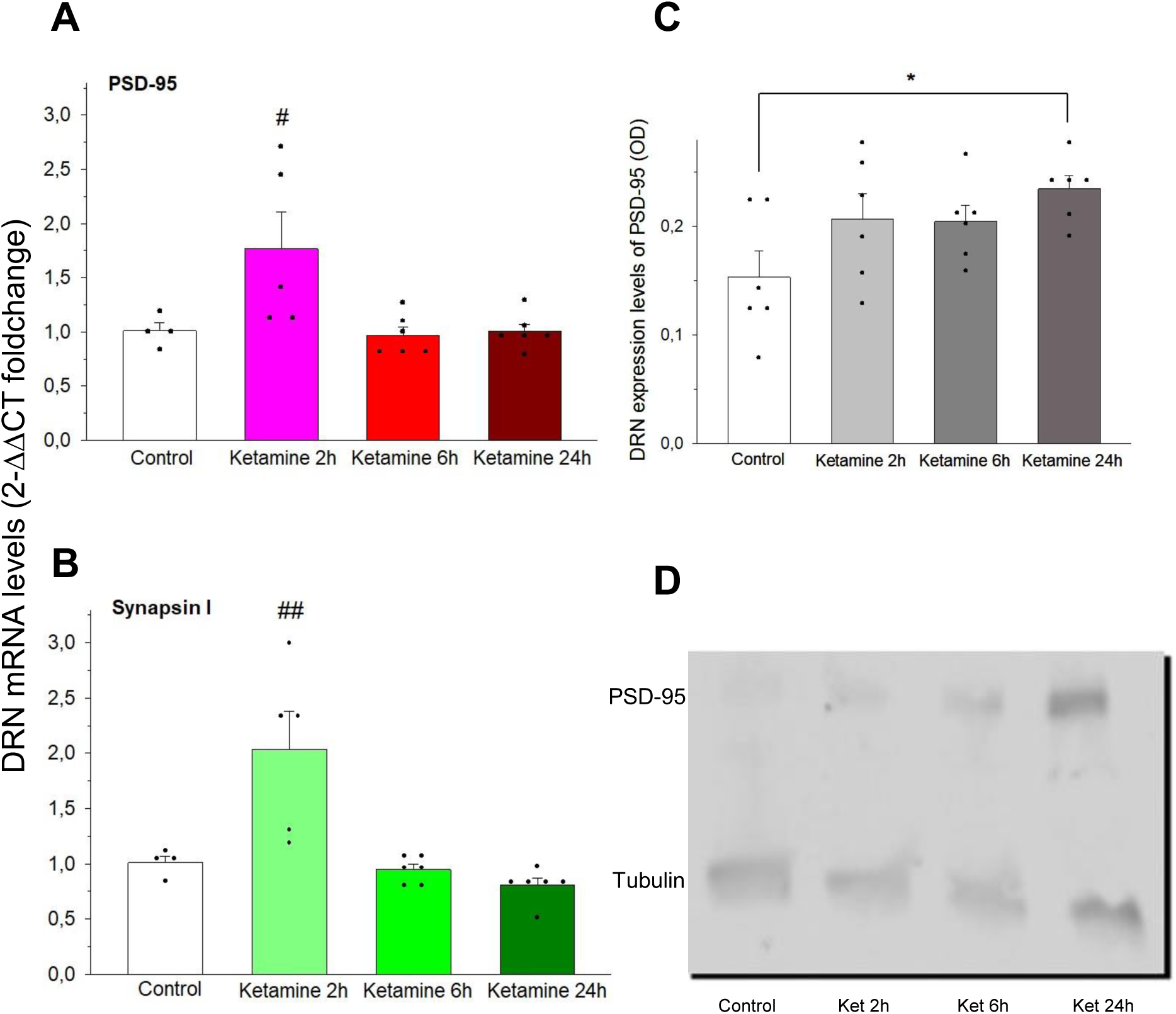
Effect of ketamine on the expression of molecular markers of synaptic plasticity within the DRN. Ketamine was injected to the animals, and the DRN micro-dissected at different time points thereafter. Control groups were obtained with samples from vehicle-treated rats, extracted two hours after the injection. **A** and **B**: Effect of ketamine administered 2, 6 or 24 hours before the micro-dissection on DRN expression of PSD-95 and synapsin I mRNA, respectively. A quantitative RT-PCR was performed, and mRNA levels compared by using the 2-ΔΔCT method, normalized with the control values. Data are expressed as mean ± SEM, each dot representing the value found in a single animal. # p < 0.05, ## p < 0.01 vs Control, Dunnett’s test after significant one-way ANOVA. **C**: Effect of ketamine administered 2, 6 or 24 hours before the micro-dissection on the protein level expression of PSD-95 within the DRN. Data are expressed as densitometric analyses of Western blots, with tubulin used as the loading control for normalization with respect to total protein levels. * p < 0.05 vs control, Student’s *t*-test. **D**: Example of an immunoblot from a typical experiment.

### Effect of ketamine and of pCPA on cell proliferation within the DG

The ability of ADs to enhance cell proliferation in the DG is considered both a prerequisite and a hallmark of their efficacy (33,43). It has previously been reported that a single subanaesthetic dose of ketamine induces such a mitogenesis already 16 h post-administration, in strong agreement with its profile of rapid AD (44). By using the thymidine analogue BrdU and the 5-HT depleter pCPA, we assessed the extent to which the mobilization of endogenous 5-HT provoked by ketamine is involved in this effect. One way ANOVAs revealed group effects, when performed either on the entire DG [F(3, 24) = 13, p < 0.001; **Fig. 5A**], or by subdividing the hippocampus in its ventral [F(3, 24) = 22.7, p < 0.001; **Fig. 5B** and **5D**] and dorsal [F(3, 24) = 6.9, p < 0.01; **Fig. 5C** and **5E**] subparts. In each case, the vehicle/ketamine group was found significantly higher than the vehicle/vehicle one (Newman-Keuls test, p < 0.05), thus confirming the existence of a facilitatory action of ketamine on DG cell proliferation. Interestingly, the increase was 30% for the entire hippocampus (**Fig. 5A**), very close to the value of 25% reported by authors using a different index of mitogenesis (44). However, this effect was abolished in the presence of pCPA, as the pCPA/vehicle and pCPA/ketamine groups did not differ one from each other, the hippocampus being taken as its whole or subdivided (Newman-Keuls tests, n.s.; **Figs. 5A-C**). These results indicate that the facilitation of dentate cell proliferation seen after the injection of ketamine is completely dependent on endogenous 5-HT. Of note, regardless of the presence of ketamine, values in pCPA-treated animals were lower than in the vehicle/vehicle ones when considering the entire hippocampus (Newman-Keuls test, p < 0.05 for both pCPA/vehicle and pCPA/ketamine; **Fig. 5A**). This effect was strong in the ventral DG (Newman-Keuls test, p < 0.001 both; **Fig. 5B**), but did not reach statistical significance in its dorsal part (Newman-Keuls tests, n.s.; **Fig. 5C**).

**Figure 5.**
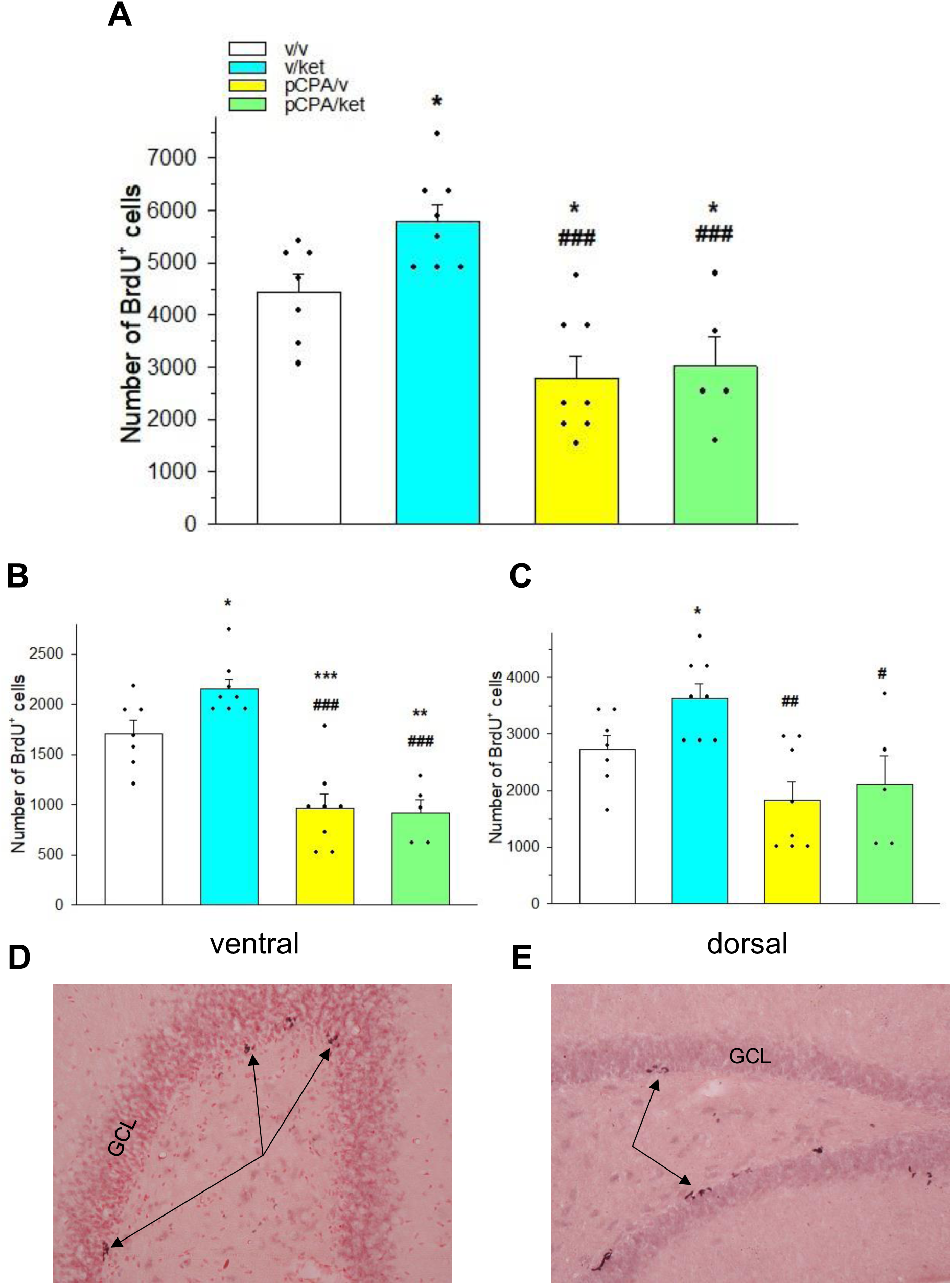
Effect of ketamine and of para-chlorophenylalanine on hippocampal cell proliferation. The thymidine analogue BrdU (50 mg/kg, i.p.) was delivered twice daily during two days, and the brain extracted 24 h after the last injection; the 5-HT depleter para-chlorophenylalanine (pCPA) or its vehicle (v) was administered once daily (150 m/kg, i.p.) 72, 48 and 24 h before the first dose of BrdU. A subanaesthetic dose of ketamine (ket; 10 mg/kg, i.p.) or of its vehicle (v) was also given to the animals the first day of BrdU administration. **A**: Number of BrdU-positive cells found in the DG, estimated on the basis of a serial counting made on every sixth section of the entire hippocampus. Data are expressed as mean ± SEM, each dot representing the value calculated in a single animal. **B** and **C**: Same results as those in panel **A**, subdivided between the ventral and dorsal parts of the hippocampus, respectively. * p < 0.05, ** p < 0.01, *** p < 0.001 vs v/v; # p < 0.05, ## p < 0.01, ### p < 0.001 vs v/ket, Newman-Keuls test after significant one-way ANOVA. **D** and **E**: Representative photomicrographs of the dentate gyrus (DG) in the ventral and dorsal hippocampus, respectively. The *arrows* point some examples of BrdU-positive cell clusters. GCL: granular cell layer. Magnification: x40.

## DISCUSSION

The present study shows that ketamine used at an AD-effective, subanaesthetic dose exerts an excitatory influence on the activity of DRN 5-HT neurons. This effect is not immediate, requiring a few hours to manifest, as it seems to depend upon a process of progressive synaptic strengthening within the DRN. This involvement of the 5-HT system appears in turn to be essential for triggering the mechanisms of neuroplasticity in the DG that coincide with the onset of a sustained AD action.

It is really relevant to consider definitions and nomenclature with caution, when discussing the kinetical features of ketamine AD properties. As reminded in the Introduction, the question has been addressed in the literature by referring to distinct temporal scales across studies. We deemed “immediate” those effects that take place within 2 hours, a duration corresponding to ketamine half-life (45), where others refer to the term “very rapid” (44). On the other hand, we chose to call “rapid” the events which occur some 24 to 48 h after ketamine administration, because this corresponds to a much faster onset than what is observed with classical AD such as SSRIs (7). However, these rapid events are also maintained over time, occurring after ketamine half-life has elapsed and being still observable up to 1-2 weeks after its administration both in clinical use and in animal models (1,46–47). For this reason, they are most usually deemed “sustained” in the relevant studies (i.e. 22,24-26,48). To put it clearly, it is implicit that the effects that we place herein in the “rapid” category are likewise “sustained”.

In agreement with earlier data (28–29), the firing rate of DRN 5-HT neurons remains unaltered up to two hours after the administration of 10 mg/kg, i.p. of ketamine. These results suggest that the 5-HT system is not involved in the immediate time scale of ketamine properties. Consistently, depleting brain 5-HT contents with pCPA does not affect the ability of ketamine to reduce immobility in the Forced Swimming Test (FST) 1 h after ketamine (22), but abolishes it when done 24 h afterwards (22,26). Not surprisingly, brain circuitries underlying the immediate and rapid/sustained effects of ketamine in the FST are distinct, the latter appearing to involve neurons originating from the ventral hippocampus, unlike the former (38). Several reports have shown that subanaesthetic doses of ketamine may increase the release of 5-HT within minutes post-administration in the mPFC of both rats and mice (20,26,48–52). Interestingly, it seems that this property results from the activation of cholinergic neurons projecting from the Pedunculopontine Tegmental Nucleus toward the DRN, which in turn recruit a subpopulation of 5-HT cell bodies (52). Our results clearly suggest that this subpopulation is small in numbers, as its involvement is not accompanied by a significant enhancement of the average 5-HT neuronal firing rate. It remains possible however that, in a brain environment hypersensitive to cholinergic transmission, such as in the Flinders sensitive rat line (53), the influence of this subpopulation becomes more prominent. This could contribute to explain why the immediate effects of ketamine (15 mg/kg, i.p.) in the FST have been reported to be dependent on the mobilization of 5-HT neurons in this specific strain (24–25).

At difference with the preceding, the 5-HT system appears to be massively involved in the AD-like properties of low-dose ketamine, with regard to what we have defined as rapid kinetics. Thus, the increase of swimming behavior observed in the FST 24 h after the drug, is suppressed by a pretreatment with pCPA both in naïve rats and in a model of stress-induced depression (22). The same result has been observed in mice, in which in addition, a positive correlation between the time spent swimming and brain 5-HT extracellular levels has been established (26). In the present study, a strong enhancement of DRN 5-HT function manifests itself after a single dose of 10 g/kg of ketamine, provided a minimum delay of ≈ 2 hours is allowed to elapse. A previous study conducted in mice had already reported that about 50% of DRN 5-HT cell bodies express the early gene cFos 90 to 120 min after the administration of ketamine (30 mg/kg, i.p.) (23). This important mobilization of 5-HT neurons is in strong contrast with the immediate effect that ketamine elicits on mPFC 5-HT efflux, which clearly recruits only a marginal amount of them (see above). The average firing rate of 5-HT cells 2-5 h post-injection was found to be 90% above control values, which is more robust than the 50-60% increase induced by other pharmacological agents known to facilitate DRN 5-HT electrical activity, such as 5-HT_4_ agonists and antagonists of metabotropic glutamatergic type 2 (mGluR2) receptors (36,54–55). These latter compounds have been shown to act onto 5-HT neurons through mechanisms involving excitatory projections issued from the mPFC (23,36,54). Similarly, we show here that the delayed positive control of ketamine is abolished when this brain region is lesioned, confirming once again the central role it plays on the regulation of central 5-HT function (31,56–58). The influence exerted in the mPFC by ketamine at low dosage has features that are quite specific, to the point that they have been characterized as “long-term potentiation (LTP)-like” (35). It is notably manifested by the synthesis of several synaptic signaling proteins which takes place within hours, and results in a progressively increased synaptogenesis in pyramidal cells (11,13). The mechanisms involved seem to be complex, putting into play both a direct action on these cells, and an indirect one *via* the blockade of NMDA receptors in GABAergic cortical interneurons (35,59–60). Importantly, the mTOR-mediated transduction pathway is triggered early in the process (1 h after ketamine) (61), of which it appears to constitute an essential actor (11,13). In our hands, a pre-treatment with the potent mTOR inhibitor Torin 2 suppressed the facilitatory effect of ketamine on 5-HT firing, suggesting that the occurrence of the LTP-like mechanism(s) in the mPFC are a prerequisite for its expression. It has already been shown that a high-frequency stimulation of this cortical region triggers the expression of pro-plastic factors in DRN 5-HT neurons (32,57). It can therefore be hypothesized that the administration of ketamine at a subanaesthetic dose provokes the emergence of a “plastic cascade” that arises in the mPFC, and then propagates in at least one of its anatomical targets, namely the DRN.

In favor of this hypothesis, we observed a progressive increase of the postsynaptic strengthening factor PSD-95 in the hours that followed the injection. The amounts of protein were still enhanced after 24 h, at difference with the transcription of its mRNA which took place in a transient manner, suggesting that the underlying processes are highly regulated at the nuclear level. Interestingly, a previous microdialysis study had reported that in the rat DRN, ketamine at 10 mg/kg does not affect the release of 5-HT up to two hours after its administration, but induces a gradual mobilization of glutamate, which extracellular levels became doubled at that time (20). On this basis, the authors proposed that the “rapid” (i.e. immediate according to our nomenclature) AD action of ketamine might be related to an efflux of glutamate, whereas the 5-HT system would be responsible for a more sustained effect of the drug (20). The delayed enhancement of 5-HT neuron firing unveiled in the present study appears indeed to have functional consequences downstream of the DRN, some of them being potentially long-lasting. Thus, in parallel with its effect on PSD-95, ketamine at 10 mg/kg increased the transcription of the mRNA coding for Synapsin I, a presynaptic factor essential to sustain the vesicular release of a neurotransmitter in response to enhanced neuronal activity (41). Kinetics of mRNA transcriptions were very similar for both the pre-and the postsynaptic marker, suggesting that the 5-HT system actually holds a pivotal role in the transfer of neuroplasticity evoked above. Consistent with this idea, our microdialysis experiments showed that in the ventral hippocampus, one of the main areas of projection for DRN 5-HT neurons (27), the release of the transmitter increased progressively starting from 2 h after ketamine administration. Obviously, the hippocampal formation also occurs to constitute a key area, regarding the therapeutical potential of treatments with AD purposes. It has been known for two decades now that chronic ADs enhance neurogenesis in the DG, and that disrupting this effect blocks their behavioral properties (33,62). Importantly, a single administration of a 5-HT agonist is sufficient to trigger DG cell proliferation within 4 hours, and with long-lasting consequences since it results in a significantly increased neurogenesis 4 weeks later (63). The present results show that low-dose ketamine also induces a rapid (2 days) mitogenesis in this brain area, an ability that appears to be fully dependent on the integrity of the central 5-HT transmission. Based on those, it can be proposed that the route taken by ketamine to produce its rapid AD-like action corresponds to a “neuroplastic wave” which originates in the mPFC, then spreads into DRN 5-HT neurons to ultimately provoke the required neurogenic changes of the DG. More recently, a bundle of data conducted to hypothesize that neurogenesis as such would be essential for the long-term sustainment of an AD treatment, but that rapid AD-like efficacy would rather result from a synaptic reinforcement of granular cells within the DG (43,64–65). In this regard, ketamine at the dose of 5 mg/kg, i.p. has been shown to restore hippocampal LTP deficits in the Wistar-Kyoto rat model of depression (66). More importantly, and in striking similarity with our present results, this effect was observed 3.5 h after the drug injection, but not when it had been administered only 30 min before the induction of LTP (66).

Altogether, the above data strongly suggest that the 5-HT system occupies a central role for the expression of the rapid AD-like properties of low-dose ketamine. Although the latter have been shown to be durable over time (46), they eventually fade away: the available clinical data indicate that they generally disappear within one week after a single injection (3). Interestingly, in rats the mean firing rate of DRN 5-HT neurons is no more modified by 10 mg/kg of ketamine, when measured 24 h after the administration (67). The facilitation of 5-HT activity evidenced in the present study appears therefore to be transient, its expression taking place in a narrow timeframe, consisting of a few hours. Even when given repeatedly (e.g. thrice in a week), ketamine has no visible effect on 5-HT neuronal activity 24 h after the last injection (67). Several hypotheses can be proposed to account for this peculiar feature. Firstly, the function of 5-HT_1A_ autoreceptors has been reported to be highly responsive to changes in DRN 5-HT neuron homeostasis (68–69). The data available so far indicate that their sensitivity is not modified ≈2 h post-ketamine (20), but further studies are needed to assess whether this is still the case at 24 h. Indeed, the present results suggest that pro-neuroplastic proteins such as PSD-95 are still expressed at this timepoint, and it cannot be excluded that the 5-HT_1A_-mediated inhibitory feedback could then be enhanced to compensate. Second, a paradoxical reduction of 5-HT neuron firing rate can occur in the presence of two excitatory treatments, likely through a phenomenon similar to a “depolarization block” (34). Because the action of ketamine on 5-HT function is prolonged, this may also play a role when the drug is administered over several days. Whatever the exact mechanisms involved, the facilitatory control exerted by low-dose ketamine on 5-HT activity appears to be pulsatile in nature. Even if single “pulses” of 5-HT transmission may be sufficient to trigger long-lasting effects in limbic areas such as the DG (see above), it will be required to repeat them across time if the objective is to sustain a therapeutical efficacy over several months, as it is the case in clinical use. A recent clinical trial has reported that some treatment-resistant depressed patients met the response criteria when ketamine was first given thrice weekly for 6 weeks (priming phase), and once a week thereafter (stabilization) (5). We hope that the present findings may help to further refine such protocols, by highlighting the relevance of optimizing the mobilization pace of the 5-HT system. One limitation of our study however, is that it did not address the influence of subanaesthetic ketamine on noradrenergic and dopaminergic neurons, which are known to contribute to AD efficacy even though their influence on DG neuroplasticity is not as prominent. These two aminergic systems actually respond to the drug with kinetics suggesting that their plasticity is modulated differently than that of 5-HT cells (67).

In summary, this study contributes to bring the importance of 5-HT in ketamine rapid AD-like action back at the center of the game, by unveiling the existence of an original mechanism that involves successive transfers of neuroplastic processes across different brain regions. It will be worth of interest in the future to assess whether similar phenomena are susceptible to occur with other pharmacological agents, either glutamatergic or not. Beyond a potential interest for clinical application, these findings may thus open new avenues of research in the neuropharmacology of the monoaminergic nuclei.

## MATERIALS AND METHODS

### Animals

Experiments were carried out in male Sprague-Dawley rats (Charles River, L’Arbresle, France and Janvier Labs, Le Genest-Saint-Isle, France) weighing 270–420 g and kept under standard laboratory conditions (12:12 light-dark cycle with free access to food and water, light starting from 7 AM). Animal experiments were performed in compliance with the European Communities Council for the care and use of laboratory animals (86/609 ECC, still available at the time the study was initiated), and with the approval of the Animal Care Committee of the University of Bordeaux and of the French Ministry of Higher Education and Research.

### Chemicals, drugs and pharmacological treatments

The following pharmacological compounds were used: ketamine (Ketamine 1000, Virbac, Carros, France), para-chlorophenylalanine (pCPA) methyl ester hydrochloride, 5-bromo-2′-desoxyuridine (BrdU) and 9-(6-Aminopyridin-3-yl)-1-(3-(trifluoromethyl)phenyl)benzo[h][1,6]naphthyridin-2(1H)-one (Torin 2) (Sigma-Aldrich, St-Quentin Fallavier, France). The solution of Ketamine 1000 (100 mg/ml) was freshly diluted 1:10 in NaCl 0.9% before each use, to reach a concentration of 10 mg/ml; a volume of 1 ml/kg was then administered to the animals. Torin 2 was diluted in a solution of β-cyclodextrin 30% (w/w) containing 10% DMSO, and the other compounds in NaCl 0.9%. For experiments aimed at measuring hippocampal cell proliferation, the tryptophan hydroxylase inhibitor pCPA was administered once daily at the dose of 150 mg/kg, i.p. during three consecutive days. The first dose of BrdU was then administered one day after the last pCPA injection (see below for details). In each set of experiments, drug dosages refer to the free base. All other chemicals and reagents were the purest commercially available.

### Extracellular recordings of DRN 5-HT neurons

Recordings were performed using single-barreled glass micropipettes. Electrodes were filled with a 0.5 M sodium acetate solution containing 2% Pontamine Sky Blue, resulting in an impedance of 10-15 MΩ. Rats were anaesthetized with chloral hydrate (400 mg/kg, i.p.) and placed in a stereotaxic frame. A burr hole was drilled on the midline 1 mm anterior to lambda. DRN 5-HT neurons were encountered over a distance of 1 mm starting immediately below the ventral border of the Sylvius aqueduct (70). Recordings were performed by using a compact, all-in-one complete electrophysiology system (ARIS-22, Neurostar GmbH, Sindelfingen, Germany) and data were analyzed with the Spike2 software (Cambridge Electronic Design), so that the firing rate was calculated as the mean number of events occurring within a 10 s period. The presumed 5-HT neurons were identified using the classical criteria: a slow (0.5-2.5 Hz) and regular firing rate and long-duration (0.8-1.2 ms) positive action potentials (34,71). In the experiments aimed at assessing the effect of ketamine alone in naïve or lesioned (see below) rats, ketamine or its vehicle (NaCl 0.9%) was administered i.p. at the start of the surgery, so that recordings began after a minimal amount of 2 hours after the injection. Several “descents” were made along the DRN in each animal to record 7 to 12 neurons and this, until up to 5 hours after ketamine. In the experiments using the Torin 2 compound (see below), recordings started 30 min after the administration of ketamine, and lasted up to 4 hours thereafter. Additional chloral hydrate doses (50 mg/kg, i.p.) were given when necessary to maintain anaesthesia.

### Involvement of the mPFC: pre-treatment with Torin 2 and electrolytic lesions of the mPFC

Torin 2 (1 mg/kg, i.p.) or its vehicle was administered 15 min before ketamine, thus 45 min before the recordings started. Electrolytical lesions of the mPFC were performed immediately before starting the surgical procedure for DRN extracellular recordings, as previously described (34). A stainless-steel electrode (0.3 mm diameter) was inserted bilaterally into the mPFC at the following coordinates: AP = 3.4 mm, L = ± 0.5 mm with respect to bregma, H = 4.6 mm from the surface of the skull (70). A constant DC source was used to pass a 500 µA current through the electrode for each side, for a duration of 10 s. The electrode was removed at the end of the lesion procedure.

### Measurement of hippocampal 5-HT extracellular levels by intracerebral microdialysis

Rats were anaesthetized with 3% isoflurane, and placed in a stereotaxic frame equipped with a stereotaxic anaesthesia mask. A microdialysis probe (CMA/11, cuprophan, 4 mm long, 240 μm outer diameter, Carnegie Medicin, Phymep, France) was implanted in the right ventral hippocampus (coordinates, in mm: AP = −5.4, L = 4.9 relative to bregma, H = −7.8 relative to the cortical surface). After the surgery, the percentage of isoflurane was adjusted to 1.2% until the end of the experiment. Probes were perfused at a constant flow rate (0.5 μl/min), by means of a microperfusion pump (CMA 111, Carnegie Medicin, Phymep), with aCSF containing (in mM): 147 NaCl, 4 KCl, 2.2 CaCl2, pH 7.4. Pharmacological treatments were performed 120 min after the beginning of the perfusion (stabilization period). 5-HT outflow was monitored during 240 min after the injection of ketamine. Dialysates fractions were collected in a refrigerated fraction collector (MAB 85 Microbiotech, Phymep) every 20 min. At the end of each experiment, the animal was deeply anesthetized with a pentobarbital overdose (400 mg/kg, i.p.), and its brain was removed and fixed in NaCl (0.9%)/paraformaldehyde solution (10%). Probe location was determined histologically on serial coronal sections (60 μm) stained with cresyl violet, and only data obtained from rats with correctly implanted canulae were included in the results.

After collection, dialysate samples were immediately analyzed with a high-performance liquid chromatography apparatus (Alexys UHPLC/ECD Neurotransmitter Analyzer, Antec, The Netherlands), equipped with an autosampler (AS 110 UHPLC cool 6-PV, Antec), as previously described (72). The mobile phase [containing (in mM) 100 phosphoric acid, 100 citric acid, 0.1 EDTA.2H2O, 1.1 octanesulfonic acid. NaCl plus 6% acetonitrile, adjusted to pH 6.0 with NaOH solution (50%)] was delivered at 0.070 ml/min flow rate with a LC 110S pump (Antec) through an Acquity UPLC BEH column (C18; 1 × 100 mm, particle size 1.7 μm; Waters, Saint-Quentin-en-Yvelynes, France). Detection of 5-HT was carried out with an electrochemical detector (DECADE II, Antec) with a VT-03 glassy carbon electrode (Antec) set at +460 mV *versus* Ag/AgCl. Output signals were recorded on a computer (Clarity, Antec). Under these conditions, the retention time for 5-HT was 4–4.5 min. and the sensitivity was 50 pM with a signal/noise ratio of 3:1. 5-HT content in each sample was expressed as the percentage of the average baseline level calculated from the three fractions preceding drug administration. Data correspond to the mean ± S.E.M. of the percentage obtained in each experimental group.

### Quantification of PSD-95 and Synapsin I mRNA levels by quantitative RT-PCR

Rats were treated with a single dose of ketamine (10 mg/kg, i.p.) or its vehicle, and sacrificed by decapitation 2, 6 and 24 hours thereafter. Their brains were removed, and the region corresponding to the DRN was micro dissected for mRNA quantification. Trizol extraction of total RNA was carried out according to the manufacturer specifications (Ambion). Thus, 0.5 ml of Trizol was added to each sample; following incubation for 5 min, 0,1ml of chloroform was then added. This mixture was centrifuged at 14,000 × g for 15 min at 4 °C, before the addition of 0,25ml of Isopropanol to the aqueous phase. These samples were then mixed and centrifuged again at 14,000 × g for 25 min. The supernatant was discarded, the RNA pellet was washed with 1 ml of 75% ethanol, centrifuged at 7,500 × g for 5 min, and air-dried in sterile conditions for 5–10 min. The RNA pellets were dissolved in 50 μl/20μl of RNase-free water and stored at −80 °C. Total DNA was digested with Turbo DNase from Ambion, according to the manufacturer instruction (Ambion).

Total RNA was then processed and analyzed following an adaptation of previously published methods (73). cDNA was synthesized from 2μg of total RNA using RevertAid Premium Reverse Transcriptase (Fermentas) and primed with oligo-dT primers (Fermentas) and random primers (Fermentas). QPCR was perfomed using a LightCycler® 480 Real-Time PCR System (Roche, Meylan, France). QPCR reactions were done in duplicate for each sample, using transcript-specific primers, cDNA (4ng) and LightCycler 480 SYBR Green I Master (Roche) in a final volume of 10μl. The PCR data were exported and analyzed in an informatics tool (Gene Expression Analysis Software Environment) developed at the NeuroCentre Magendie (Bordeaux, France). For the determination of the reference gene, the Genorm method was used (74). Relative expression analysis was then normalized against two reference genes: for these experiments, the peptidylprolyl isomerase a (Ppia) and ribosomal protein L13a (Rpl13a) ones were selected. The relative level of expression was calculated using the comparative (2-ΔΔCT) method (75). For PSD-95 (Gene: Dlg4, GenBank ID: NM_019621), the primer sequences were CTTCTCAGCCATCGTAGAGG (forward) and GAGAGGTCTTCAATGACACG (reverse); for Synapsin I (Gene: Syn1, GenBank ID: NM_019133), they were ACCCCTCTCTCCTGGTTCTACAT (forward) and AGGGACATCTTGAGGAATGAGAAG (reverse).

### Western blot analyses

Rats were treated with a single dose of ketamine (10 mg/kg, i.p.) or its vehicle, and sacrificed by decapitation 2, 6 and 24 hours thereafter. Their brains were removed, and the region corresponding to the DRN was micro dissected and homogenized in boiling 1% SDS. Protein concentrations were measured, in duplicate, using DC-protein assay (Biorad, Hercules, CA). Protein extracts (25μg) were separated by SDS-PAGE tris-glycine gels (Invitrogen, Burlington, CA) and transferred to nitrocellulose membranes (Invitrogen, Burlington, CA). Blots were incubated with the primary antibody overnight at 4°C (mouse anti-PSD-95, 1:1000; Santa Cruz biotechnology). After several washing, sections were exposed (1 h) to the secondary antibody (anti-mouse, 1/10000, CST) in blocking buffer. Proteins were detected with the ECL plus detection reagents (Amersham Biosciences) using an LAS-3000 imaging system (Fujifilm, Düsseldorf, Germany). Relative intensities of the labelled bands were analyzed by densitometric scanning using the ImageJ software. For quantification, tubulin was used as a loading control for the evaluation of total protein levels (mouse anti-tubulin, 1/2000, Sigma).

### Evaluation of Hippocampal Cell Proliferation

For analysis of BrdU-positive cells, rats received 4 injections of BrdU (50mg/kg i.p.) or its vehicle (two injections per day spaced 6h apart, two consecutive days). On day 1, animals also received a single i.p. injection of ketamine (10 mg/kg, i.p.). Twenty-four hours after the last BrdU injection (thus 48 h after ketamine), rats were deeply anesthetized with pentobarbital (400mg/kg, i.p.) and transcardially perfused (150mL of 0.9% NaCl followed by 500mL of 4% paraformaldehyde). After perfusion, all brains were post-fixed overnight in paraformaldehyde at 4°C and stored at 4°C in 30% sucrose. Coronal sections (30 µm-thick) were cut through the entire hippocampus, between −1.60 to −6.72, from Bregma (70) on a freezing microtome, and stored at 4°C in 10mM phosphate-buffered saline (PBS) pH 7.4 containing 0.1% NaN3.

Free-floating sections were then incubated with 50% formamide and 50% 2x saline sodium citrate buffer (pH 7.4; 0.3M sodium citrate, 0.03M NaCl) for 90min at 65°C, and immersed in 2x saline sodium citrate buffer, followed by 30min in 2M hydrochloric acid (HCl) to denature deoxyribonucleic acid (DNA). After neutralization with 0.1M sodium borate (pH 8) and pre-incubation with PBS containing 0.3% Triton X-100 (PBST), sections were processed for 3min in pepsin (0.45U/mL in 0.1M HCl) and transferred for 15min to 3% H_2_O_2_ to eliminate endogenous peroxidases. Then, sections were incubated at room temperature overnight with a primary antibody against BrdU (rat monoclonal IgG, 1/400, AbCys) in PBST 1% bovine serum albumin. Sections were then washed with PBST and incubated with a secondary antibody for 90 min (biotinylated rabbit anti-rat IgG, 1/250, Vector Laboratories), followed by amplification with an avidin-biotinylated horseradish peroxidase complex (Elite ABC kit, Vector Laboratories). Peroxidase activity was revealed by incubating sections with 0.02% 3,3′-diaminobenzidine and 0.8% NiCl in 0.5mol/L Tris-HCl (pH 7.4). After several rinses, sections were mounted on gelatine-coated slices, dehydrated in graded alcohols, and cover-slipped in DePeX mounting medium (VWR).

The number of BrdU-labeled cells was quantified with a light microscope (Zeiss) at 40x magnification on a series of every sixth section of the entire hippocampus, by an experimenter unaware of the treatments underwent by the animal. The hippocampus was segregated into two halves, with the dorsal hippocampus defined as the anterior region (−1.60 to −5.30 Bregma) and the ventral hippocampus defined as the posterior region (−5.30 to −6.72 Bregma). About 10 sections in the dorsal hippocampus and 5 sections in the ventral hippocampus were examined per animal to estimate the total number of BrdU-labeled cells per structure. BrdU-labeled cells were counted in the granule cell layer and the subgranular zone, and cells that were located more than two cells away from the subgranular zone were omitted.

### Statistical analysis

All data are expressed as means ± SEM unless otherwise specified. In electrophysiological recordings, the average firing rate was established for each animal, on the basis of 7-12 neurons recorded in successive tracks performed along the DRN. These average values were then used for statistical assessment. In the experiments aimed at studying the effects of ketamine in the presence of pCPA and Torin 2, a one-way ANOVA was used to address the presence of a group effect, followed when significant by the Newman-Keuls *post-hoc* test for multiple comparisons. The different kinetics of ketamine on the expression levels of PSD-95 and synapsin I mRNA and/or protein within the DRN were assessed by using a within-designed one-way ANOVA, followed when significant by the Dunnett’s *post-hoc* test with the control (vehicle) group taken as the reference. When necessary, a Student’s *t*-test was also applied vs the control group (see Results for details). Similarly, when ketamine was administered alone, the Student’s *t*-test was used to allow paired-wise comparisons with the control conditions. Finally, in the single case where data did not follow a normal distribution, non-parametric methods (one-way ANOVA on ranks followed by the *post-hoc* Dunn’s test) were required to assess for group differences (see Results for details).

## ACKNOWLEDGMENTS

This study was funded by grants from the Agence Nationale de la Recherche (ANR-2009-MNPS-026.01 and ANR EMMA 11-050), as well as by financial supports from the INSERM, the CNRS, the Université de Bordeaux, the Université Claude Bernard Lyon 1 and the Université de Nice-Sophia Antipolis.

## Notes

### Competing Interest Statement

The authors have declared no competing interest.

